# Plasticity of plant defense and its evolutionary implications in wild populations of *Boechera stricta*

**DOI:** 10.1101/144626

**Authors:** Maggie R. Wagner, Thomas Mitchell-Olds

**Author notes:** Correspondence and requests for materials should be addressed to M.R.W.

## Abstract

Phenotypic plasticity is thought to impact evolutionary trajectories by shifting trait values in a direction that is either favored by natural selection (“adaptive plasticity”) or disfavored (“nonadaptive” plasticity). However, it is unclear how commonly each of these types of plasticity occurs in natural populations. To answer this question, we measured glucosinolate defensive chemistry and reproductive fitness in over 1,500 individuals of the wild perennial mustard *Boechera stricta,* planted in four common gardens across central Idaho, USA. Glucosinolate profiles—including total glucosinolate quantity as well as the relative abundances and overall diversity of different compounds—were strongly plastic both among habitats and within habitats. Patterns of glucosinolate plasticity varied greatly among genotypes. More often than expected by chance, glucosinolate profiles shifted in a direction that matched the direction of natural selection, indicating that plasticity among habitats tended to increase relative fitness. In contrast, we found no evidence for within-habitat selection on glucosinolate reaction norm slopes (i.e., plasticity along a continuous environmental gradient). Together, our results indicate that glucosinolate plasticity may improve the ability of *B. stricta* populations to persist after migration to new habitats.

## Introduction

The role of phenotypic plasticity in adaptive evolution has been a subject of great controversy and research interest for decades (Bradshaw 1965; Via and Lande 1985; Via *et al.* 1995; Pigliucci 2005; Ghalambor *et al.* 2015; Hendry 2015). It has long been recognized that both an organism’s genotype and its environment shape its phenotype, which then determines its evolutionary fitness. Strictly speaking, phenotypic variation caused by environmental stimuli is not heritable and therefore cannot result in evolution through systematic changes in allele frequencies (Falconer and Mackay 1996). Nevertheless, plasticity is predicted to impact evolution by shifting phenotypes that are under natural selection (Bradshaw 1965). Furthermore, if patterns of plasticity are genetically variable, then plasticity itself may evolve in response to selection (Gomulkiewicz and Kirkpatrick 1992). It remains unclear how commonly these phenomena occur in natural populations, and whether the adaptive value of plasticity varies for different traits, environments, and spatial scales.

One way that plasticity could impact evolution is by accelerating or hindering adaptation to a novel environment— e.g., upon invasion of a new habitat or in response to a relatively sudden ecosystem shift, as might result from climate change (Donohue *et al.* 2001; Richards *et al.* 2006; Ghalambor *et al.* 2007; Anderson *et al.* 2012). Plasticity that moves a phenotype closer to the new phenotypic optimum is often called “adaptive” plasticity because it increases fitness relative to a non-plastic genotype (Figure 1g–i); however, whether this type of plasticity actually facilitates genetic adaptation is controversial. Strong adaptive plasticity could place an organism very near to the new adaptive peak, removing the selective force that would otherwise drive adaptation through allele frequency change and thereby inhibiting local adaptation. Alternatively, moderate adaptive plasticity may enable a population to survive in the new environment long enough for selection to increase the frequency of beneficial alleles, thus promoting local adaptation (Baldwin 1896; Price *et al.* 2003; Ghalambor *et al.* 2007). The opposite pattern—in which “nonadaptive” plasticity moves phenotypes farther from the new optimum—may either increase the risk of extinction or lead to rapid adaptive evolution by intensifying natural selection (Conover and Schultz 1995; Ghalambor *et al.* 2007; Ghalambor *et al.* 2015; Huang and Agrawal 2016). In this manuscript, we do not attempt to determine whether plasticity constrains or facilitates long-term genetic adaptation. Rather, our goal is to assess the relative frequency of adaptive versus nonadaptive plasticity in natural populations.

**Figure l.**
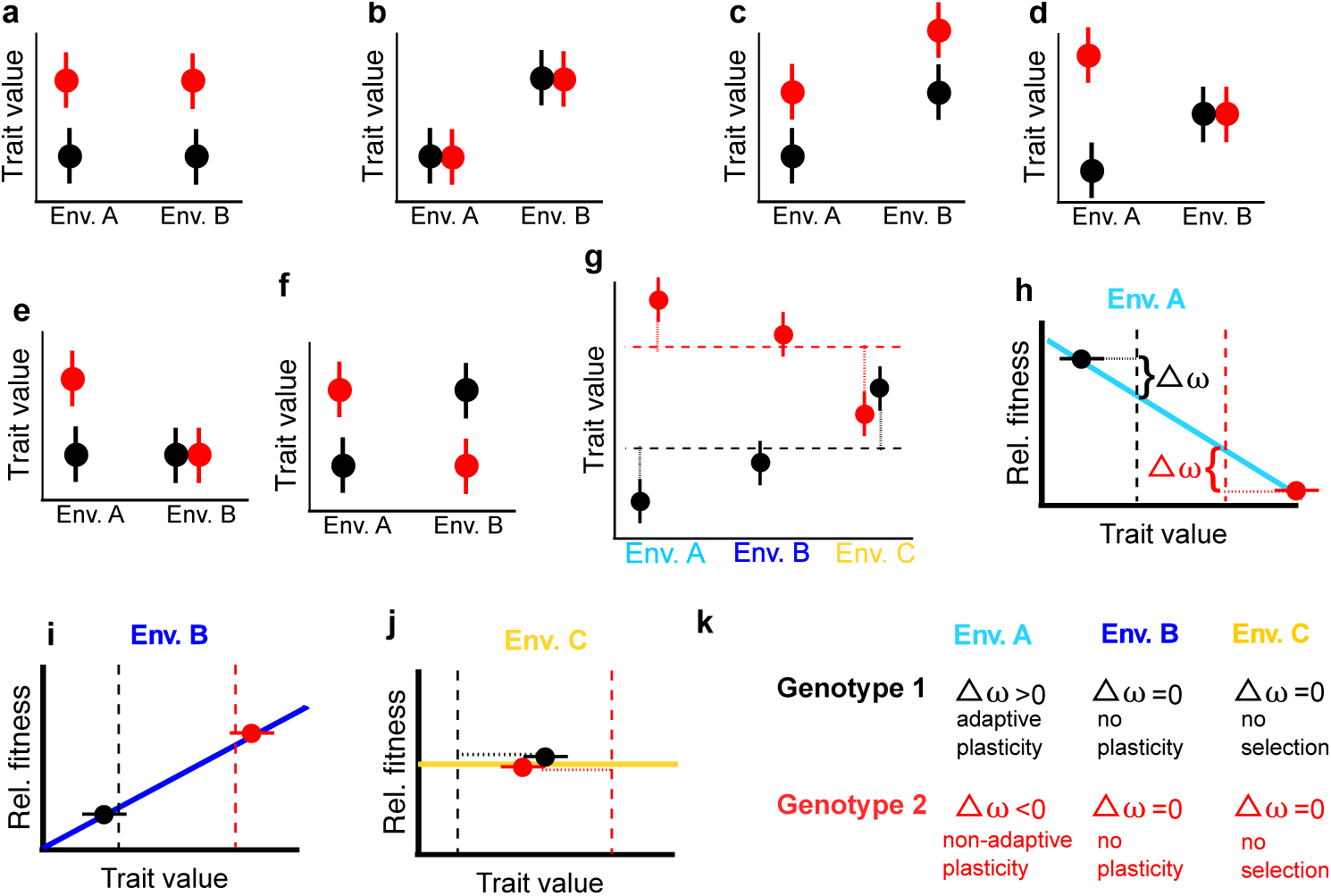
Schematic of genotype-by-environment interactions and adaptive plasticity between discrete environments. Points show mean trait values of two genotypes (“red” and “black”). Panel **(a)** depicts a trait that is under pure genetic control with no plasticity. Panel **(b)** depicts a trait that is plastic but not genetically variable. Panel **(c)** shows both a genotype effect and plasticity, but no interaction between them. Panels **(d)-(f)** depict examples of genotype-by-environment interactions. In **(d),** a genetic difference is detectable only in one site; the genotypes have plasticity of equal magnitude but opposite sign so that the mean phenotype in each site is identical. In **(e),** only one genotype is plastic. In **(f),** the genotypes switch rank phenotype; averaged across sites, there is no genetic difference between them, and the average trait value within each site is the same. Note that panels **(a)-(f)** could represent six different traits measured simultaneously in one experiment: plasticity is a property of a particular trait, a particular genotype, and a particular environmental change. Panels **(g)-(k)** illustrate how “adaptive” and “nonadaptive” plasticity among environments can be detected. First, each genotype’s trait value in each site can be compared to its average trait value across environments (depicted as dashed red or black lines, horizontal in **g** and vertical in **h-j,** representing a hypothetical genotype that is identical except that it is completely nonplastic). Next, a selection differential (an equation expressing relative fitness as a function of trait values, shown as solid colored lines in **h-j)** is calculated for each environment. To determine the expected change in fitness due to plasticity, the selection differential is evaluated at the “no-plasticity” baseline trait value (dashed lines) and at the genotype’s true trait value observed in that environment. **(k)** In hypothetical Environment A, one genotype shows adaptive plasticity and the other shows nonadaptive plasticity. In Environment B, neither genotype shows a significant plastic deviation from its experiment-wide mean. In Environment C, both genotypes show substantial plastic deviations, but the trait is not under selection so the plasticity has no effect on fitness.

A related but distinct question is whether, and how, plasticity might evolve as an adaptation to environmental heterogeneity within a single habitat.

Plasticity in response to fine-scale environmental variation is often imagined as a *reaction norm,* with the trait value as some function of a continuous environmental predictor (Figure 2; Schmalhausen 1949). For traits that exhibit genotype-by-environment interactions, genetic variation exists for reaction norm shape. Natural selection acting on variation for plasticity can be detected using established quantitative genetic methods such as genotypic selection analysis on reaction norm coefficients (Figure 2g–h; Weis and Gorman 1990; Rausher 1992; Baythavong and Stanton 2010). Natural selection is predicted to result in increased plasticity if patterns of plasticity are heritable, if the spatial scale of changing selective pressures is similar to the organism’s dispersal distance (Levins 1962; Gomulkiewicz and Kirkpatrick 1992; Baythavong 2011), if reliable environmental cues for the selection pressure are available (Levins 1963; Donohue *et al.* 2000; Schmitt *et al.* 2003; Reed *et al.* 2010), and if costs of plasticity are minimal (Auld *et al.* 2010).

**Figure 2.**
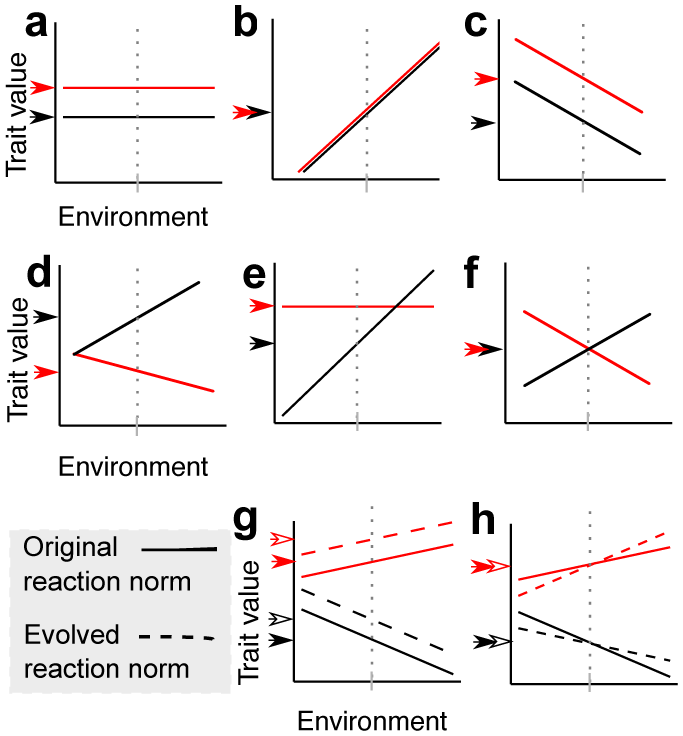
Genotype-environment interactions and selection on plasticity over continuous environmental gradients: Examples of reaction norms. Lines show the mean trait values of two genotypes (“red” and “black”) across a range of some continuous environmental predictor. The arrows on the vertical axis indicate reaction norm *height,* or the trait value for each genotype evaluated at the average value of the environmental predictor (indicated by grey tick marks on horizontal axis and vertical dotted lines). Panel **(a)** shows a genetic difference with no plasticity— i.e., zero slope. Panel **(b)** shows plasticity with no genetic difference. Panel **(c)** shows both plasticity and a genotype effect, but no interaction between them, indicated by parallel reaction norms—the genotypes differ in reaction norm height, but not slope. Panels **(d)-(f)** all show possible genotype-by-environment interactions, or genetic variation for reaction norm shape. In panel **(d)** reaction norm slopes differ in sign but not magnitude; in **(e)** only one genotype is plastic. In panel **(f)** the genotypes are indistinguishable when averaged across all environments (e.g., if the environmental gradient was unobserved); the genotype difference is environment-dependent. When mean trait values and plasticity are genetically uncorrelated, they can evolve independently. Panel **(g)** illustrates changes in reaction norm height in response to linear selection for increased mean trait values. Panel (h) illustrates evolutionary change in reaction norm slope in response to linear selection for more positive linear reaction norm coefficients. Note that such selection may result in either increased or decreased overall plasticity (steeper or shallower slope), depending on the original shape of the reaction norm.

Despite several excellent empirical studies (Dudley and Schmitt 1996; Schmitt *et al.* 1999; Donohue *et al.* 2000; Donohue *et al.* 2001; Sultan 2001; Baythavong 2011), more examples from natural populations are needed to test theoretical predictions about the fitness impacts of both between-environment and within-environment plasticity (Hendry 2015). Data on the plasticity and evolution of physiological traits (as opposed to morphological or life-history traits) is particularly scarce (Palacio-López *et al.* 2015). Because variation in phytochemistry may affect not only the evolution of the plant but also entire communities and ecosystems (Wimp *et al.* 2007; Hopkins *et al.* 2009), phytochemical plasticity has been identified as a high priority research target (Hendry 2015). Here, we address these needs by studying plasticity and evolution of glucosinolate defensive chemistry in the wild perennial herb *Boechera stricta*, a close relative of Arabidopsis. Goals of this study were (1) to characterize genotype-by-environment interactions underlying glucosinolate variation in *B. stricta*, (2) to assess whether glucosinolate plasticity alters relative fitness after transition to novel habitats, and (3) to test whether natural selection acts on glucosinolate reaction norms within habitats.

We measured glucosinolate profiles, size, and fecundity of 25 *B. stricta* genotypes replicated in 80 experimental blocks divided among four common gardens in diverse habitats (Figure 3). Because *Boechera* has limited dispersal (<0.5 m on average; Bloom *et al*. 2002), the environmental variation encompassed by the widely separated common gardens is much greater than what individual *B. stricta* populations normally encounter; thus, plasticity among field sites describes plasticity after a sudden environmental change or migration to a new habitat. To assess whether glucosinolate plasticity among habitats exhibits an “adaptive” or “nonadaptive” pattern, we compared the direction of selection in each site with the direction of diverse genotypes’ plastic responses to that site (Figure 1g–k). Then, we quantified within-habitat glucosinolate plasticity and assessed its relationship to fecundity in each habitat using genotypic selection analysis on reaction norm coefficients (Figure 2g–h). We found that substantial genotype-by-environment interactions underlie glucosinolate variation in *B. stricta*, and plasticity among sites tended to move trait values in an adaptive direction; however, we did not detect selection on glucosinolate plasticity within habitats.

**Figure 3.**
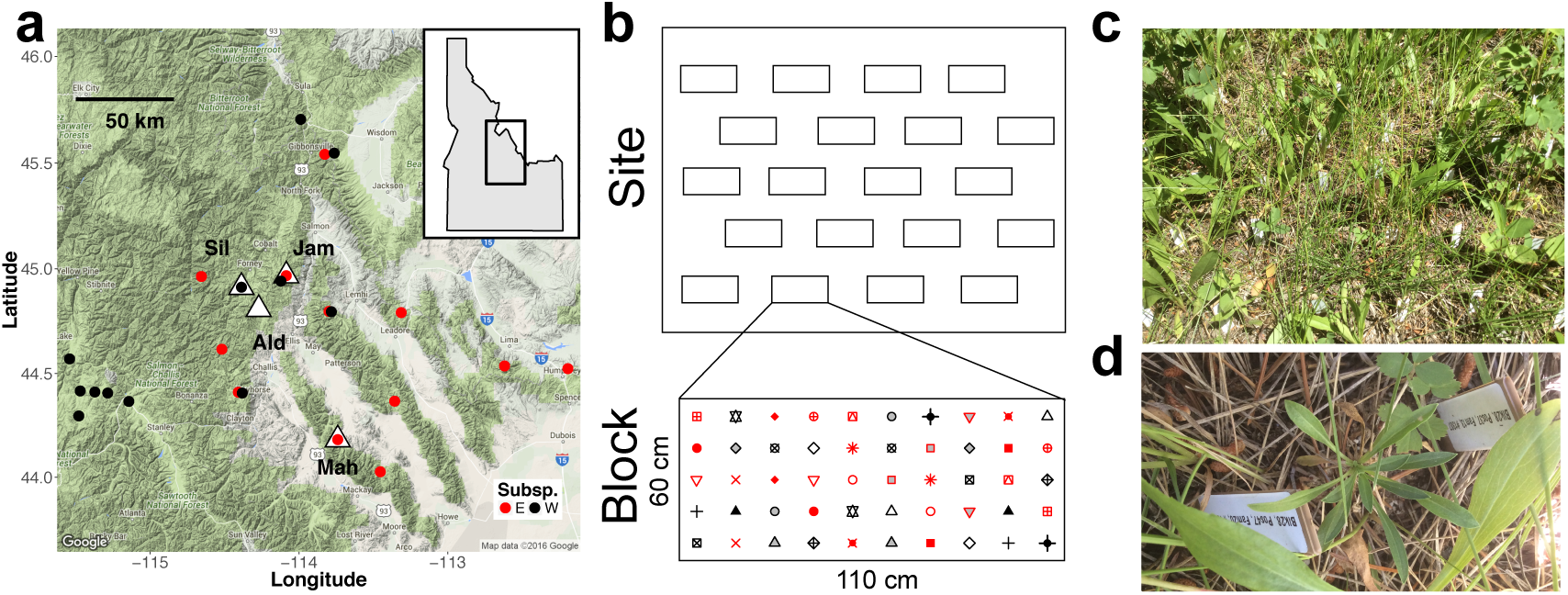
Field sites and experimental design. **(a)** Map of field sites and wild *B. stricta* populations used in common garden experiment. Common gardens are denoted with white triangles and labeled. Circles mark collection sites of the 25 genotypes included in the experiment. Not shown: one EASTERN genotype collected in Colorado. Map data: Google. **(b)** Each common garden contained 16 to 22 experimental blocks. Each randomized block contained two individuals of each of the 25 genotypes, planted in a 10-cm grid. Panel **(c)** shows a block placed within the natural vegetation; **(d)** shows one experimental rosette with its identifying tag.

## Methods

All statistical analyses were performed in R version 3.3.2 (R Core Team 2016) with heavy use of the packages ggplot2, lme4, lmerTest, dplyr, tidyr, and stringr (Wickham 2009; Bates *et al.* 2015; Kuznetsova 2015; Wickham and Francois 2015; Wickham 2016a, b). Throughout, *P*-values were corrected for multiple comparisons using the sequential Bonferroni correction (Holm 1979). Additional details for all sections are available in Supplementary Methods. All data and R code will be made freely available in a Dryad repository upon publication.

### Study system

The short-lived perennial herb *Boechera stricta* (Graham) Al-Shehbaz is common in montane meadows and forests throughout its native range in WESTERN North America (Rushworth *et al*. 2011). Natural populations are strongly genetically differentiated (*F*_ST_=0.56; Song et al. 2006) and have adapted to diverse habitats that vary in climate, water availability, elevation, soil composition, plant community diversity and density, and microbial community composition (Supplementary Figure 1; Anderson *et al.* 2013a, b; Wagner *et al.* 2016).

*B. stricta* produces a variety of glucosinolates, which are sulfur-rich, biologically active phytochemicals that protect against generalist insect herbivores and pathogens and may also affect nonpathogenic root-associated microbes (Agrawal 2000; Tierens *et al*. 2001; Brader *et al*. 2006; Halkier and Gershenzon 2006; Bednarek *et al.* 2009; Bressan *et al.* 2009; Hopkins *et al.* 2009; Schranz *et al.* 2009; Sanchez-Vallet *et al.* 2010). Glucosinolates are constitutively produced, although attack by natural enemies often induces additional production (Agrawal 1998, 2000; Brader *et al.* 2001; Agrawal *et al.* 2002; Textor and Gershenzon 2009; Abdel-Farid *et al.* 2010; Manzaneda *et al.* 2010). *B. stricta* produces four aliphatic glucosinolate compounds with varying biological activity (Figure 4a; Windsor *et al.* 2005; Schranz *et al.* 2009; Prasad *et al.* 2012). Total glucosinolate concentration and relative abundances of these compounds vary extensively among individuals (Figure 4b–c).

**Figure 4.**
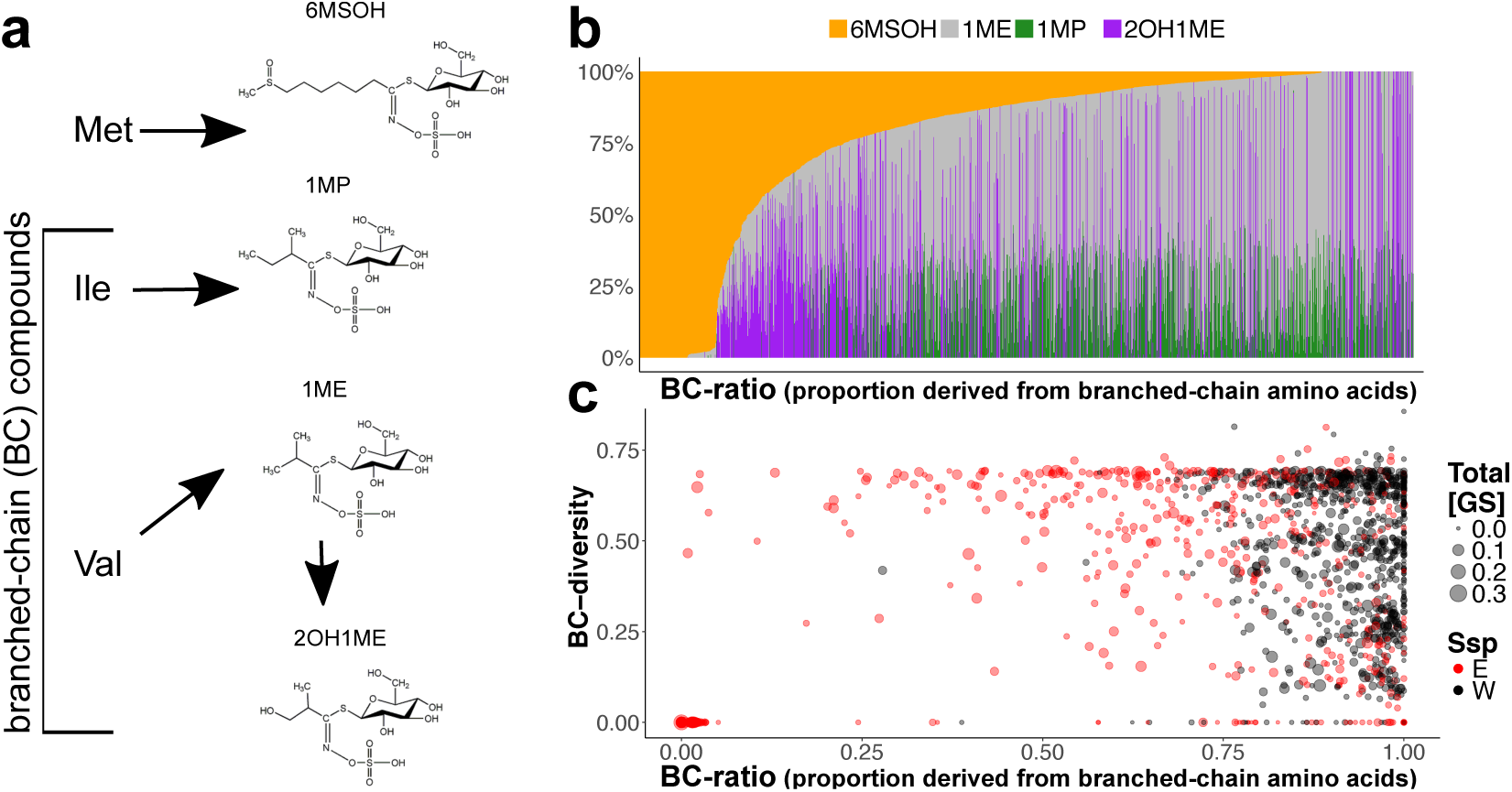
Glucosinolate variation in 1,505 field-grown *Boechera stricta* rosettes. **(a)** Chemical structures of the four primary glucosinolates in *B. stricta* (Kanehisa *et al.* 2002), with their amino acid precursors. 6MSOH is derived from methionine; the other compounds are all derived from branched-chain amino acids. **(b-c)** Rosette leaf glucosinolate profiles of 1,505 field-grown plants. In both panels, plants are sorted by increasing BC-ratio (i.e., decreasing 6MSOH). **(b)** The proportions of four aliphatic glucosinolates are shown for each measured plant. **(c)** Three summary metrics of each glucosinolate profile are shown for each individual. Three genotypes in this study lack branched-chain glucosinolate functionality and only produce 6MSOH; individuals of these genotypes are seen in the lower left-hand corner of panel **(c).**

### Design and installation of field experiment

In October 2013, we planted 4,000 self-full siblings of 25 naturally inbred *Boechera stricta* genotypes (Supplementary Table 1) in fully randomized blocks (two replicates per genotype per block) in four common gardens in central Idaho (Figure 3). These field sites are all home to wild *B. stricta* populations, and are distinguished by many biotic and abiotic environmental characteristics (Supplementary Table 2; Supplementary Figure 1). Each genotype was derived from an accession from one wild *B. stricta* population (Figure 3a), which we propagated by self-fertilization in standard greenhouse conditions to minimize variation caused by maternal environmental effects. These genotypes represent the breadth of *B. stricta* genetic diversity, comprising 12 from the WEST subspecies and 13 from the EAST subspecies (Lee and Mitchell-Olds 2013). Because *B. stricta* primarily self-pollinates and is naturally inbred (*F*_IS_=0.89; Song *et al.* 2006), self-full siblings are essentially genetically identical. Therefore, phenotypic variation among individuals of the same genotype describe that genotype’s plastic response to environmental variation.

### Measurement of plant performance in the field

During summer 2014, we returned to each site several times to measure survival, developmental stage, and height. At the end of the growing season, we measured fruit production for each surviving individual. Because *B. stricta* is predominantly self-pollinating (Song *et al.* 2006), fruit production reflects both male and female fecundity, and thus is a good estimate of reproductive fitness.

For phenotypic selection analyses (below), we used fecundity (in mm of fruit produced) as a measurement of reproductive fitness:

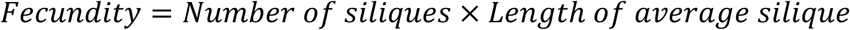

For estimation of genotypic fitness, we also calculated the probability of survival for each genotype *l*:

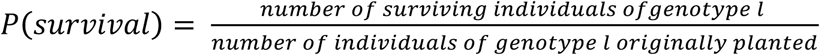

We then calculated the total evolutionary fitness for each genotype as:

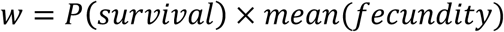

### Measurement of glucosinolate profiles

Because insect attack can induce additional production of glucosinolates (Agrawal 1998), we measured glucosinolate profile as early as possible in the summer, before peak herbivory. On the earliest census date for each site, we collected ~20-30 mg of rosette leaf tissue from each surviving plant into tubes containing 70% methanol. Samples were shipped to Duke University, then fully randomized onto 96-well plates. Glucosinolates were extracted from the methanol leachates using established protocols (Supplementary Methods). We used high-performance liquid chromatography (HPLC) to measure the abundance of four aliphatic glucosinolates (Figure 4a–b) in each sample. Three of these compounds (1ME, 1MP, 2OH1ME) have branched-chain structures and biological activity that differs from that of the fourth, straight-chain compound (6MSOH; Figure 4a; Schranz *et al.* 2009; Prasad *et al.* 2012). We calculated absolute concentrations (*μ*mol per mg dry weight) of each compound by comparing each peak to an internal standard and dividing by the dry weight of each leaf sample (Supplementary Methods).

From the absolute concentrations of all four compounds, we calculated three summary metrics for each sample’s glucosinolate profile:

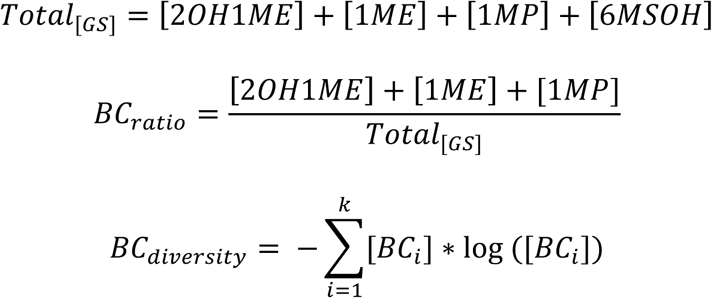

where *k* = the total number of branched-chain compounds present in the sample and [*BC_i_*] = the concentration of the *i*^th^ branched-chain glucosinolate. Total [GS] describes the combined concentration of all aliphatic glucosinolates. BC-ratio describes the proportion of aliphatic glucosinolates that are derived from branched-chain amino acids, which is an ecologically and evolutionarily important trait in *B. stricta* (Schranz *et al.* 2009; Manzaneda *et al*. 2010; Prasad *et al.* 2012). Finally, BC-diversity describes the balance of the three types of branched-chain glucosinolates, taking low values when glucosinolate profiles are dominated by one compound and high values when multiple compounds are present in similar amounts (Figure 4b–c). We calculated BC-diversity using the Shannon diversity index in the **R** package **vegan** (Oksanen *et al.* 2013).

### Partitioning variance in glucosinolate profiles

To assess plasticity of glucosinolate profiles among habitats, we used univariate REML mixed models and ANCOVA to partition variance in each of the three glucosinolate traits among genetic and environmental predictors. We modeled each trait as:

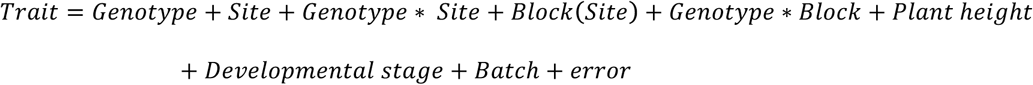

where *Height*, *Developmental Stage*, and *Batch* were nuisance variables to control for (respectively) the “general vigor problem” of large plants having more resources to invest in defense (Agrawal 2011), ontogenetic changes in rosette glucosinolate profiles, and HPLC batch effects. *Block* (nested in *Site*), *Genotype*\**Block*, and *Batch* were random-intercept terms; the rest were modeled as fixed effects. One genotype was omitted from the analysis because no individuals of that genotype survived at one field site. Spearman’s rank correlation tests of least-squares means resulting from this model (for the Genotype fixed effect) revealed that these traits were partially genetically correlated, although correlations between BC-ratio and both other traits were driven by three outlier genotypes that entirely lacked branched-chain glucosinolate functionality (Supplementary Table 3). Least-squares mean trait values for *Site*, *Genotype,* and *Genotype* × *Site* fixed effects were used to quantify plasticity among habitats.

To better understand how variation in individual glucosinolate compounds underlies variation in these three emergent properties of glucosinolate profiles, we repeated the above analysis for square-root-transformed concentrations of 2OH1ME, 1ME, 1MP, and 6MSOH. Similar to the emergent properties that are the focus of this study, concentrations of individual compounds were strongly genetically controlled but also highly plastic within and among habitats. (Supplementary Tables 4, 9b; Supplementary Figures 2-3). However, genetic correlations between individual compounds were even stronger than those between emergent glucosinolate profile properties (Supplementary Table 3b). For this reason, and because previous work has confirmed the ecological and evolutionary importance of emergent glucosinolate profile properties (Prasad et al. 2012; Müller et al. 2010), we focus the remainder of the analyses on BC-ratio, Total [GS], and BC-diversity rather than individual compounds.

### Characterizing selection on glucosinolate profiles

We evaluated natural selection on three separate glucosinolate-related traits (Figure 4c) in each of four common gardens (Figure 3) by conducting twelve phenotypic selection analyses (3 glucosinolate traits × 4 sites). The within-site relative fecundity of all measured individuals was regressed onto standardized trait values, while controlling for microsite variation in habitat quality using a random intercept *Block* term:

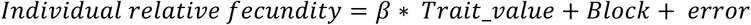

A significant *β* regression coefficient indicated nonzero linear selection. This regression coefficient is the selection differential on one trait at one site (*e.g.*, the slopes of the lines in Figure 1h–j). The signs and magnitudes of the regression coefficients from these models thus indicate the direction and strength of linear selection on each trait at each site.

Next, to test whether selection differentials for each trait varied among sites (i.e., whether selection was spatially variable), we analyzed the relative fecundity and standardized phenotype data from all four sites together by fitting a mixed-effects ANCOVA model with an additional *Site* × *Trait* interaction term, which described heterogeneity of linear selection on the trait among habitats:

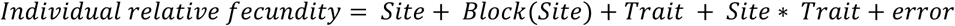

Because stabilizing or disruptive selection could also affect the fitness consequences of plasticity (i.e., movement of trait values either toward or away from optimum values at each site), we also conducted quadratic selection analysis for each trait at each site:

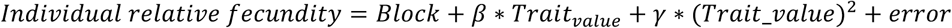

A significant *γ* regression coefficient indicated nonzero quadratic selection; positive values were taken as evidence of stabilizing selection, while negative values indicated disruptive selection. However, quadratic regressions were included in downstream analyses only if they gave significantly improved fit over a linear selection model, and if we could reject the null hypothesis that the true fitness minimum or maximum fell outside the range of observed trait values (Mitchell-Olds and Shaw 1987).

### Testing for adaptive plasticity among habitats

The question of whether plasticity can aid survival in new environments hinges on whether the plastic response moves trait values in a direction favored by natural selection, thus increasing relative fitness in the new environment. We explored this interaction between plasticity and selection by combining results from the variance-partitioning model and selection analyses described above. We restricted this analysis to combinations of sites and traits where we detected significant linear or quadratic selection.

For each site-trait combination, we calculated each genotype’s expected change in fitness due to plasticity (Δ*ω*) by evaluating the selection differential at two trait values:

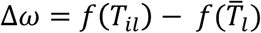

where *f*(*x*) is the selection differential (i.e., relative fitness as a function of a given trait at a given site; see above), *T_il_* is the least-squares mean trait value of genotype *l* at site *i* (depicted by the black and red points in Figure 1g–j), and *T*̅*_l_* is the least-squares mean trait value of genotype *l* averaged across all sites (depicted by the black and red dashed lines in Figure 1g–j). Both *T_il_* and *T*̅*_l_* were calculated from the REML variance-partitioning model described above. *T*̅*_l_* represents the trait value of a hypothetical genotype that is identical to genotype *l* except that it lacks plasticity. Therefore, Δ*ω* is an estimate of the fitness change that can be attributed to plasticity among sites of a trait that is under selection. Positive values of Δ*ω* constituted evidence of adaptive plasticity; negative values indicated non-adaptive plasticity.

We conducted an exact binomial test of the null hypothesis that plasticity is equally likely to move glucosinolate trait values in an “adaptive” or a “non-adaptive” direction. Weak cases of plasticity, in which the 95% CI of Δ*ω* included zero, were excluded from this analysis.

### Characterizing within-habitat plasticity using reaction norms

In this study, we focused on phenotypic plasticity induced by spatial environmental variation at a single time-point. Because each plant in this study only experienced a single spatial environment, plasticity of individual plants could not be measured. Instead, spatial plasticity of glucosinolate profiles is a property of a genotype, estimated by comparing the phenotypes of individuals that shared the same genotype but were growing in different experimental blocks. To infer whether natural selection was acting on fine-grained glucosinolate plasticity within *B. stricta* habitats, we (1) quantified plasticity among blocks for each genotype as a continuous function or *reaction norm*, and (2) used genotypic selection analysis to test whether reaction norm steepness—a measure of plasticity—predicted evolutionary fitness of each *B. stricta* genotype.

First, for each of 25 genotypes we fit one reaction norm to describe each of the three glucosinolate traits as a continuous linear function of an *environmental index* (EI, a numerical descriptor of microhabitat conditions within each experimental block)—for a total of 75 reaction norms (3 traits × 25 genotypes). Data from all four sites were pooled for calculation of reaction norms. We assumed that most environmental factors causing glucosinolate plasticity are unknown, and so the relevant environmental characteristics are best “measured” using plant phenotype data. Therefore, we assigned the grand mean trait values observed in each block (for all 25 genotypes, pooled) to be the environmental indices (Finlay and Wilkinson 1963). We then calculated a reaction norm for each genotype and each trait using linear regression of genotype-specific block mean trait values onto the EIs:

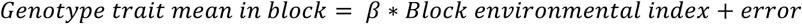

The linear regression coefficient (*β*) estimated by each model is a reaction norm *slope*, which describes the magnitude and direction of the plastic response for one genotype. We also calculated the *height* of each reaction norm by evaluating the linear function at the mean EI value; thus, reaction norm height describes the genotype’s predicted trait value in an “average” block (Figure 2).

Second, to test whether reaction norm slopes were heterogeneous among genotypes, we analyzed genotype-specific block mean trait values and EIs from all 25 genotypes together by fitting an ANCOVA model with an additional *Genotype* × *EI* interaction term that described genetic variation for reaction norm slopes:

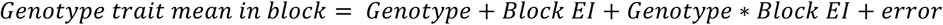

If the interaction term was significant, we concluded that reaction norm slopes were heterogeneous among genotypes.

### Testing for selection on reaction norm slope and reaction norm height

The slope and height of the reaction norms are measurable characteristics of plant genotypes, corresponding to plasticity and average trait values, respectively (Figure 2). We measured linear selection gradients on these reaction norm parameters to determine whether glucosinolate plasticity in response to fine-grained environmental variation affects fitness. Reaction norm parameters were calculated using data pooled from all four sites (see above), but to allow for the possibility that plasticity is not equally advantageous in all habitats, we measured selection on these parameters separately at each site.

To test for linear selection on reaction norm slopes, we conducted genotypic selection analysis separately for each glucosinolate trait at each site. Genotypes’ relative fitness within each site (genotypic survival rate x genotypic fecundity, divided by the mean fitness for all genotypes at that site) was regressed onto the genotypes’ reaction norm height and slope, generating two partial regression coefficients (*β*1 and *β*2) that describe linear selection on average trait values and trait plasticity, respectively (Figure 2g–h; Supplementary Methods):

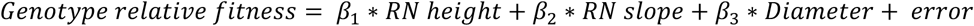

The mean rosette diameter for each genotype at time of transplanting, which reflects differences in performance in the greenhouse and not in the field, was included as a nuisance variable.

## Results

### Genotype and environment synergistically controlled glucosinolate profiles

All three glucosinolate profile features—BC-ratio, Total [GS], and BC-diversity—varied among genotypes and among field sites (Figure 5a; Table 1). All three traits were also affected by developmental stage and plant size (Table 1; Supplementary Figure 4). Genotypic variation for BC-ratio has been previously reported in *B. stricta* (Schranz et al. 2009; Manzaneda et al. 2010); our data show that Total [GS] and BC-diversity are also genetically controlled. In this experiment the EAST subspecies harbored more genetic variation for BC-ratio than the WEST; however, genetic variability for the other traits was comparable between the subspecies (Figure 5a). Genotypes varied less in glucosinolate quantity than in BC-ratio and BC-diversity, except for SAD12, which produced nearly twice the concentration of glucosinolates as the others. SAD12 is also notable as the only genotype in the experiment that originated in Colorado; the others are from Idaho or Montana. Nevertheless, analysis of Total [GS] yielded the same results regardless of whether the outlier SAD12 was included.

**Figure 5:**
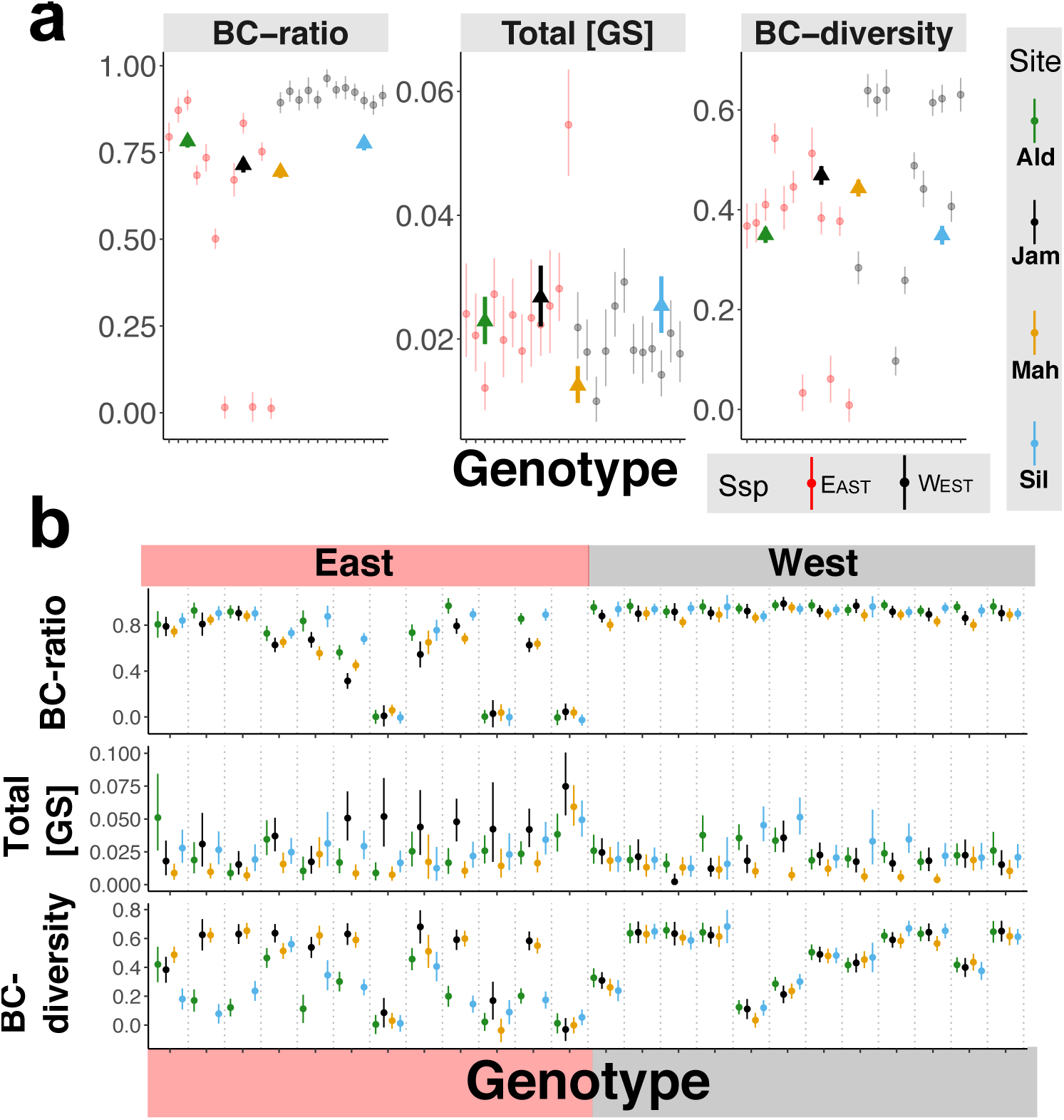
Genotype and environment influence glucosinolate profiles both independently and synergistically. **(a)** Both genotype and habitat influence glucosinolate profiles. Plotted are least-squares mean trait values for each Genotype (circles) and for each Site (triangles) from a REML mixed model that also controlled for developmental stage, plant size, genotype-by-site interactions, and block and batch effects (Table 1). Thus, circles show the mean trait value for each genotype (averaged across all sites); triangles show the mean trait value at each site (average for all genotypes). Note that the horizontal position of the triangles is meaningless—they were placed in order to not obscure the genotype means. Error bars are 95% confidence intervals. **(b)** Glucosinolate plasticity among sites is genetically variable. Genotypes are delimited by vertical dashed lines. The points are least-squares mean trait values for each genotype in each common garden, with 95% confidence intervals. BC-ratio and BC-diversity are unitless; units for Total [GS] are μmol/mg dry tissue.

**Table 1.** Statistics from REML mixed models of glucosinolate traits reveal strong genotype-by-site interactions. All effects are fixed except for Block, Genotype × Block, and HPLC batch, which were random-intercept terms. Significance of random effects was assessed using likelihood ratio tests. *N*=1,503 for Total [GS]. For BC-ratio and BC-diversity, *N*=1,486 because these traits could not be measured for individuals that produced no glucosinolates.

|  | **BC-ratio** | **Total [GS]** | **BC-diversity** |
| --- | --- | --- | --- |
| $R^2$ | 0.99 | 0.49 | 0.92 |
| Genotype | $F_{23,1035}=343.68$<br>$P < 3e^{-16}$ | $F_{23,949}=9.24$<br>$P < 3e^{-16}$ | $F_{23,982}=140.98$<br>$P < 3e^{-16}$ |
| Site | $F_{3,109}=21.28$<br>$P = 1.3e^{-10}$ | $F_{3,99}=14.07$<br>$P = 1.0e^{-07}$ | $F_{3,1085}=49.28$<br>$P < 3e^{-16}$ |
| Genotype × Site | $F_{69,1035}=3.93$<br>$P < 3e^{-16}$ | $F_{69,899}=3.02$<br>$P = 7.2e^{-14}$ | $F_{69,972}=12.91$<br>$P < 3e^{-16}$ |
| Dev. Stage | $F_{2,785}=4.31$<br>$P = 0.014$ | $F_{2,1332}=23.78$<br>$P = 2.1e^{-10}$ | $F_{2,1226}=6.49$<br>$P = 0.0031$ |
| Height | $F_{1,1109}=33.05$<br>$P = 2.3e^{-08}$ | $F_{1,1344}=46.86$<br>$P = 3.5e^{-11}$ | $F_{1,1328}=6.60$<br>$P = 0.010$ |
| Block | $\chi^2_1=22.74$<br>$P = 3.7e^{-06}$ | $\chi^2_1=49.23$<br>$P = 4.2e^{-14}$ | $\chi^2_1=4.3e^{-11}$<br>$P=1$ |
| Genotype × Block | $\chi^2_1=58.90$<br>$P = 5.0e^{-14}$ | $\chi^2_1=0.03$<br>$P=0.86$ | $\chi^2_1=12.37$<br>$P = 0.00087$ |
| HPLC batch | $\chi^2_1=4.09$<br>$P = 0.043$ | $\chi^2_1=9.66$<br>$P = 0.006$ | $\chi^2_1=8.33$<br>$P = 0.0078$ |

All three traits showed significant plasticity among sites. For Total [GS], the magnitude of the site effect was comparable to variation attributed to genotype—in particular, plants growing at Mahogany Valley produced only 50% the quantity of glucosinolates as those growing at the other sites, on average (Figure 5a). In contrast, for BC-diversity and especially BC-ratio, the magnitude of plasticity among sites was minor compared to the variation due to genotype. Additionally, BC-ratio and Total [GS] varied significantly among blocks within sites (6.3% and 11.2% of the total variance, respectively), indicating that meter-scale environmental heterogeneity affected expression of these traits (Supplementary Figure 5a; Table 1).

Finally, significant genotype-by-site interactions confirm that genotype and environment acted synergistically to shape glucosinolate profiles (Table 1). In general, EASTERN genotypes were more sensitive to environment than WESTERN genotypes, especially for BC-ratio and BC-diversity. However, patterns of plasticity among sites varied even within subspecies (Figure 5b). In addition to these genotype × site interactions, genotype × block interactions accounted for 65.3% and 35.9% of the variance in BC-ratio and BC-diversity, respectively. This indicates strong genetic variation for plasticity of glucosinolate composition in response to environmental heterogeneity on the meter scale within habitats. Consistent with the observed subspecies difference in plasticity *among* habitats (Figure 5b), EASTERN genotypes displayed greater plasticity among blocks *within* habitats, as well (Supplementary Figure 5b).

### Evidence for frequent adaptive plasticity among sites

#### Selection on glucosinolate profiles in all habitats

Phenotypic selection analysis revealed seven cases of linear selection on glucosinolate traits, and two cases of disruptive (quadratic) selection (Figure 6; Supplementary Table 5). High Total [GS] was associated with decreased fecundity in all four common gardens (Figure 6b–e; Supplementary Table 5). Pooling the data from all sites and testing for a Site*Total [GS] interaction failed to reject the null hypothesis that these selection differentials were equivalent (Supplementary Table 6), suggesting that high glucosinolate concentrations are equally costly or disadvantageous in all four habitats. In contrast, linear selection on BC-diversity varied among sites (Supplementary Table 6). Specifically, high BC-diversity was associated with higher fecundity at Jackass Meadow but lower fecundity at Alder Creek and Silver Creek (Figure 6f–h); however, we detected no significant selection on BC-diversity at Mahogany Valley (Supplementary Table

**Figure 6.**
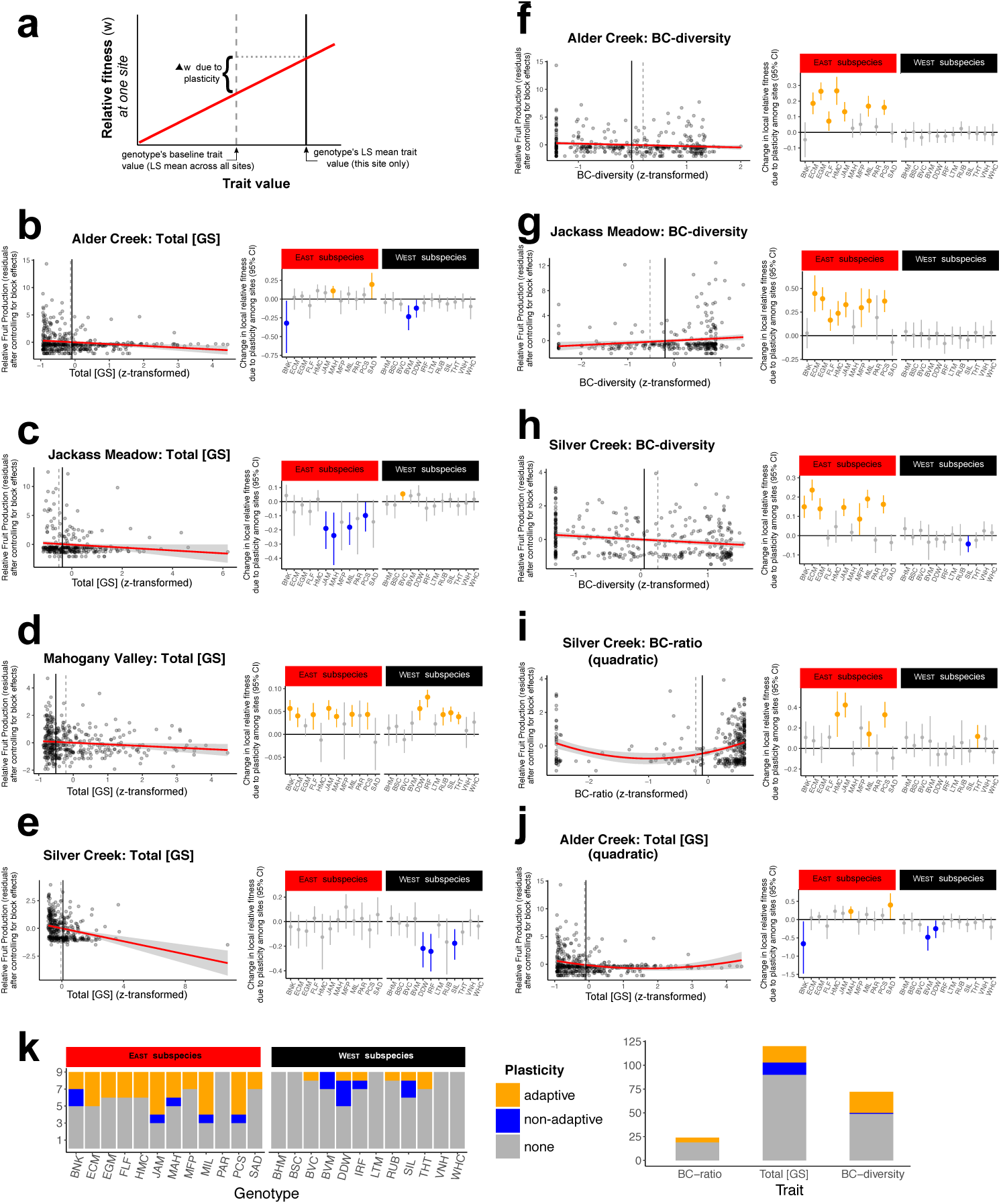
Evidence for adaptive glucosinolate plasticity among field sites. We detected seven cases in which a glucosinolate trait was under linear selection within a field site **(b-h),** and two cases of quadratic (disruptive) selection **(i-j).** Selection differentials are plotted as red lines. For each of these cases, we asked whether patterns of plasticity among sites resulted in increased or decreased relative fitness (Δ*ω*). To do this for each genotype, we evaluated the selection differential at two values: the genotype’s LS mean trait value across all sites (which represents a hypothetical “no plasticity” scenario) and the genotype’s true LS mean trait value at that site **(a).** The averages of these trait values across all genotypes are depicted as dotted and solid vertical lines, respectively, in the left side of panels **(b-h).** In the right side of panels **(b-h),** Δ*ω* due to plasticity is plotted for each genotype. Positive and negative estimates of Δ*ω* are considered evidence for *adaptive* and *non-adaptive* plasticity, respectively, if their 95% CIs do not contain 0 (colored in orange and blue respectively). Panel **(k)** summarizes counts of adaptive and non-adaptive plastic responses for each genotype (left) and each glucosinolate trait (right). Overall, plasticity was in an adaptive direction more often than would be expected by chance (44 out of 58 observations; exact binomial test, *P*=0.0001).

For thoroughness, we also calculated linear selection *gradients* to assess direct selection on each trait while controlling for indirect selection on the other glucosinolate traits (Supplementary Methods). The selection gradients generally agreed with the selection differentials, and also indicated that BC-ratio may be under negative selection at Mahogany Valley (Supplementary Table 7).

Finally, we found evidence that disruptive selection is acting on BC-ratio at Silver Creek and on Total [GS] at Alder Creek (Figure 6i–j; Supplementary Table 8).

#### Plasticity among habitats was more likely to move glucosinolate profiles in an adaptive than a non-adaptive direction

Each of the nine cases in which a glucosinolate trait was under selection at a given site (Figure 6b–j) was an opportunity for plasticity to alter trait values in a way that affected evolutionary fitness. Because the strength and direction of plasticity varied among genotypes (Figure 5), we assessed the frequency of adaptive versus non-adaptive plasticity for each genotype. We used each genotype’s experiment-wide LS mean trait values as baselines, representing a hypothetical genotype that was identical except that it lacked plasticity among sites (Figure 1g–k). Relative to this baseline, plasticity was strong enough to substantially alter relative fitness in 27% of all cases. This was especially true for genotypes of the EAST subspecies, which were more plastic than WESTERN genotypes, particularly for BC-ratio and BC-diversity (Figure 5b). Plasticity among sites affected fitness nearly 40% of the time for EASTERN genotypes, but only 14% of the time for WESTERN genotypes (Figure 6k).

Of the 58 cases in which a genotype’s plasticity was strong enough to affect its fitness, 44 shifted trait values in an adaptive direction—considerably more than would be expected by chance (exact binomial test, *P*=0.00015; Figure 6k). We found that the direction of plasticity matches the direction of selection 75.9% of the time (95% CI = 62.8% to 86.1%). However, this was largely driven by the EAST subspecies—not only because WESTERN genotypes were less plastic overall(χ^2^=13.5, *P*=0.0007), but also because their plastic responses were more likely than those of EASTERN genotypes to be in a non-adaptive direction (χ^2^=9.4, *P*=0.039; Figure 6k).

#### Glucosinolate reaction norms were genetically variable but not under detectable selection

Reaction norm slopes of all three glucosinolate traits varied strongly among genotypes, indicating substantial genetic variation for glucosinolate plasticity in response to continuous environmental gradients (Figure 7; Supplementary Table 9 a). After correction for multiple testing, genotypic selection analysis failed to detect significant selection differentials on reaction norm slopes at any site (Supplementary Table 10), suggesting that glucosinolate plasticity in response to continuous environmental gradients was neither beneficial nor costly within any of these four habitats.

### Discussion

In *Boechera stricta*, plant genotype and environment synergistically affect the concentration and composition of glucosinolate profiles. All three emergent traits of glucosinolate profiles—Total [GS], BC-ratio, and BC-diversity—were under both genetic and environmental control, with strong genotype-by-environment interactions. Genotype was the strongest determinant of BC-ratio and BC-diversity, whereas genotype and environment had similar effect sizes for Total [GS] (Figure 5a). We observed abundant genetic diversity for plasticity both among and within field sites (Figure 5b; Figure 7; Table 1). Particularly striking was the 65% of BC-ratio variation that was explained by genotype-by-block interactions (Supplementary Figure 5b; Table 1).

**Figure 7.**
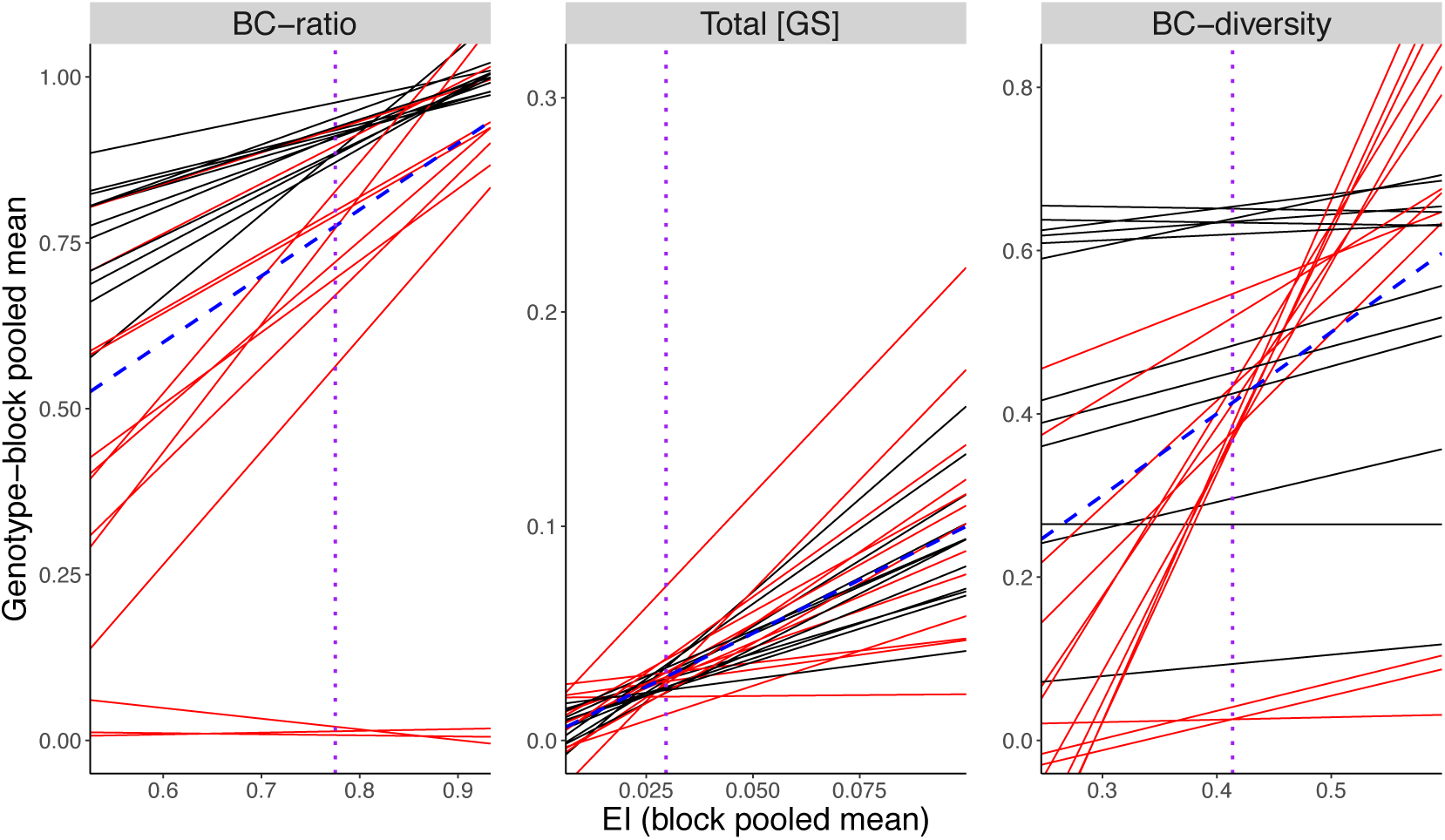
Genotype-specific glucosinolate reaction norms. The mean trait values of each genotype in each block were regressed onto the grand mean trait values for all genotypes in each block, or “environmental index”. Each resulting regression line is a reaction norm for one genotype, plotted here in red or black for EASTERN or WESTERN genotypes, respectively. Genotypes with steeper reaction norm slopes exhibit more plasticity in response to continuous environmental gradients. The “average reaction norm” (equivalent to the line *y*=1\**x*, where the expressed trait value equals the block mean trait value) is shown as a blue dashed line. The purple vertical dotted line denotes the mean environmental index for each trait (i.e., an average environment), the value at which reaction norm height was evaluated.

In general, EASTERN genotypes were more plastic than WESTERN genotypes, and there was additional genetic variation in reaction norm shape and inter-site plasticity within the EAST subspecies (Figure 5b; Figure 7). However, this divergence does not reflect adaptation to within-habitat heterogeneity. With a few exceptions, all of the environmental variables that we measured had similar variances in the two EASTERN habitats and the two WESTERN habitats (Supplementary Figure 6), indicating that neither subspecies generally occupies more complex habitats than the other. The lack of variation for glucosinolate plasticity among WESTERN genotypes is consistent with observed patterns of reduced molecular diversity relative to EASTERN genotypes (Baosheng Wang, personal communication), and might limit further evolution of reaction norms within the WEST but not the EAST subspecies.

#### Glucosinolate plasticity may aid colonization of new habitats

Because the distances separating our field sites are much greater than the dispersal distance of *Boechera* (Bloom *et al.* 2002), it is unlikely that the intersite plasticity observed in this study evolved as an adaptation to heterogeneity among sites (Via and Lande 1985; Gomulkiewicz and Kirkpatrick 1992). Instead, the differences between pairs of sites are reasonable simulations of transitions to novel environments, or when streams carry seeds to lower-elevation sites. Thus, we could assess whether this coarse-grained glucosinolate plasticity might impact the likelihood that a *B. stricta* population could survive a major environmental change or colonize a new habitat (Figure 1; Ghalambor *et al.* 2007).

Whether plasticity of a trait improves the relative fitness of a population upon encountering a new environment depends on the directions and magnitudes of both natural selection and the plastic response. Characterizing plastic responses in this way requires the definition of a “baseline” against which the site-specific trait values can be compared; here we used each genotype’s experiment-wide mean trait value as the baseline. We note that if the four field sites used in this experiment were somehow unrepresentative of the wider range of habitats occupied by wild *B. stricta* populations, then this baseline might not reflect the “true” average glucosinolate profile, and thus our estimations of plasticity might be incorrect. We have no reason to suspect this is the case, because we chose these sites to reflect the diversity of habitat types in the region (Supplementary Figure 1; Supplementary Table 2): Alder Creek and Silver Creek are riparian meadows—representative habitats of the WESTERN subspecies—while the other two sites are typical EASTERN subspecies habitats—dry, high-elevation meadows (Lee and Mitchell-Olds 2011, 2013). Nevertheless, repetition of this experiment in a wider range of *B. stricta* habitats would be necessary to rule out this possibility.

In this experiment, we detected natural selection acting on at least one glucosinolate trait at all four field sites (Figure 6b–j). Selection consistently favored lower Total [GS], suggesting a cost of producing these defensive compounds (Mauricio 1998). In contrast, the direction of selection on BC-diversity varied among sites. Strikingly, plasticity of BC-diversity in EASTERN genotypes mirrored these varying selection pressures. Trait values increased in sites where BC-diversity was under positive selection, and decreased where it was under negative selection, resulting in significant fitness boosts for the most plastic genotypes (Figure 6f–h). Across all traits, sites, and genotypes, plasticity was often not strong enough to substantially affect fitness. When it was, however, it was much more likely to increase fitness than to decrease it (Figure 6k; exact binomial test, *P*=0.00015).

The whole of the data suggest that glucosinolate plasticity often changes defensive chemistry to better match the local selection pressures, and therefore might aid *B. stricta* populations in colonizing new habitats (Ghalambor *et al.* 2007). Whether such adaptive plasticity promotes glucosinolate evolution in the long term, however, will be a more difficult question to answer (Ghalambor *et al.* 2007; Ghalambor *et al.* 2015; Huang and Agrawal 2016). Identifying the environmental cues perceived by the plants that induce changes in defensive chemistry, as well as the ecological causes of local selection pressures, should be a priority for future research on adaptive glucosinolate plasticity.

Finally, we highlight the implications of genetic variation for plasticity. Although glucosinolate plasticity was adaptive on average, some genotypes were more likely than others to exhibit nonadaptive plasticity or to simply lack plastic responses (Figure 6k). Based on our results, we expect glucosinolate plasticity to aid the colonization of new habitats 20.4% of the time, overall. However, considering only WESTERN genotypes, this estimate drops to 6.5%, while EASTERN genotypes are expected to benefit from glucosinolate plasticity 34.3% of the time. Similarly, glucosinolate plasticity is expected to *hinder* colonization of new habitats 6.5% of the time species-wide, but the rate is slightly higher for WESTERN than EASTERN genotypes (Figure 6k). These results illustrate that the contribution of glucosinolate plasticity to persistence after environmental change is not uniform across the species.

#### No evidence for selection on plasticity within habitats

In this experiment, we detected little evidence for selection on glucosinolate reaction norms, or plasticity in response to continuous environmental gradients. Because this study included only 25 genotypes, evidence for selection on reaction norms may become clearer as more genotypes are analyzed. Consistent with this, selection gradients on reaction norm height (i.e., mean trait values across all blocks) lacked statistical support but agreed qualitatively with the patterns detected using the phenotypic selection analysis (compare *β* values in Supplementary Table 5 with *β_H_* values in Supplementary Table 10). Another possible reason for this negative result is that selection pressures on glucosinolate profiles may not vary on such a fine spatial scale, reducing the opportunity for adaptive plasticity within habitats (Via and Lande 1985; Gomulkiewicz and Kirkpatrick 1992). The lack of observed selection *against* plasticity suggests that glucosinolate plasticity does not carry a significant cost (Auld *et al.* 2010).

In addition, inter-annual variation is one potential cause of plasticity and variable selection that we did not address in this study. Other experiments have shown that herbivory pressure on *B. stricta* varies considerably over a span of a few years within a single site (Mitchell-Olds, unpublished); consecutive generations of a *B. stricta* lineage might therefore experience very different predatory environments, potentially causing temporally heterogeneous selection on glucosinolate profiles. The consistent negative selection on glucosinolate quantity in this study (Figure 6b; Supplementary Table 5) suggests that herbivory intensity may have been lower than usual, reducing the usefulness of these expensive phytochemicals (Mauricio and Rausher 1997; Mauricio 1998). Temporal fluctuations in resource availability (e.g., due to limited or abundant rainfall) or pathogen pressure could also affect the relative costs and benefits of glucosinolate profile properties. Because *B. stricta* is a perennial, even a single individual may need to adjust to several environments over its lifetime; in theory, plasticity could evolve as an adaptation to such fine-scale variability (Gomulkiewicz and Kirkpatrick 1992; Reed *et al.* 2010; Baythavong 2011). The relative importance of temporal environmental variation compared to spatial variation could be assessed by extending a similar experimental design over multiple growing seasons. Follow-up studies should prioritize temporal variation as a potential driver of adaptive plasticity in glucosinolate profiles and other important *B. stricta* traits.

Importantly, evidence of natural selection on reaction norms would not be sufficient to demonstrate that plasticity evolved as an adaptation. Reliable environmental cues for the different selection pressures are also required, preventing the evolution of plasticity as an adaptation to truly stochastic environmental fluctuations (Levins 1963; Donohue *et al.* 2000; Schmitt *et al.* 2003; Reed *et al.* 2010). In other words, if an organism’s offspring reliably encounter multiple contrasting selection pressures, and if some perceptible environmental factor predicts how selection will act, plasticity (rather than habitat specialization) will evolve as a long-term adaptation to a heterogeneous environment (Donohue *et al.* 2000; Sultan and Spencer 2002). Even when these conditions are not met, however, plasticity can still improve short-term relative fitness—that is, it may sometimes be “adaptive” even if it did not evolve as an “adaptation” (Sultan 1987).

#### Future directions

The molecular basis of genotype-by-environment interactions is a key research goal for understanding the evolution of phenotypic plasticity and the robustness of genetic improvements in crop species (El-Soda *et al.* 2014). The ample genetic variation for plasticity of three glucosinolate traits provides an opportunity to explore this phenomenon at the molecular level in *B. stricta*, which offers resources such as near-isogenic lines varying in glucosinolate synthesis genes, a recently developed genome-wide association population, a fully sequenced genome, and genetic similarity to the model plant *Arabidopsis thaliana* (Rushworth *et al.* 2011; Prasad *et al.* 2012; Lee *et al.* 2017). The *Arabidopsis* glucosinolate biosynthetic pathway is well characterized (Halkier and Gershenzon 2006), although genes affecting plasticity are not necessarily part of this core pathway. Indeed, in *Arabidopsis*, patterns of flux through this pathway were robust to several biotic and abiotic environmental stimuli (Olson-Manning *et al.* 2015), suggesting that variation in environment-sensing genes or other upstream genes may be more important for glucosinolate plasticity, *per se*.

The exact ecological causes of glucosinolate plasticity in this experiment are still unclear and cannot be determined from this dataset; additional experiments with environmental manipulations will be necessary to identify the causal stimuli (Anderson *et al.* 2014). Many environmental features distinguish the field sites used in this study (Supplementary Figure 1). For this reason, we used plant phenotype to define the environmental index of each block, rather than guessing which environmental parameters were most relevant (Finlay and Wilkinson 1963). The disadvantage of this method is that the phenotype is a “black box” obscuring the actual ecological features driving trait plasticity. The question of which ecological factors drive plasticity of glucosinolate profiles—and selection on them—will be an important step towards understanding the apparent link between natural selection and phenotypic plasticity in diverse habitats across a natural landscape.

Finally, it is an open question whether the patterns of adaptive plasticity observed in this experiment are typical of other functional traits. Although glucosinolates are known to be ecologically and evolutionary important, they represent only a fraction of the traits contributing to evolutionary fitness in *B. stricta* and other Brassicaceae. Additional experiments are needed to clarify whether the evolutionary implications of glucosinolate plasticity are generalizable to other adaptively important traits in *B. stricta* and other study species. Until more examples from wild populations are available, it will remain challenging to identify patterns and draw general conclusions about the role of plasticity in adaptive evolution.

## ACKNOWLEDGEMENTS

We thank K. Donohue and M. Rausher for valuable discussion and critical comments on earlier versions of the manuscript; K. Ghattas for help in the greenhouse; R. Keith, C. Rushworth, E. Raskin, J. Lessing, and M. Olszack for help with fieldwork. M.R.W. was supported by a Graduate Research Fellowship and a Doctoral Dissertation Improvement Grant DEB-1311440 from the National Science Foundation, a Rosemary Grant Award from the Society for the Study of Evolution, and the American Philosophical Society Lewis and Clark Fund for Exploration and Field Research. T.M.-O. was supported by grant R01 GM086496 from the National Institutes of Health and EF-0723447 from the National Science Foundation.

